# Integrative Computational Analysis Reveals *H. pylori* GroEL as a Stabilizer of Neurotoxic Amyloid-β Oligomers

**DOI:** 10.1101/2025.10.12.679596

**Authors:** Wazeer Muhammed Abdullah, Ishtiaq Ahmad

## Abstract

Alzheimer’s disease (AD) is characterized by amyloid-β (Aβ) aggregation, with soluble oligomers implicated as the most neurotoxic species. Recent evidence suggests microbial infections, including *Helicobacter pylori*, contribute to AD pathogenesis. This study investigates the role of *H. pylori* GroEL, a conserved chaperonin found in bacterial outer membrane vesicles (OMVs), in stabilizing toxic Aβ oligomers. A pan-genome analysis of 353 *H. pylori* strains identified GroEL as a highly conserved protein present in 83% of strains, which supported its widespread relevance. We structurally modelled a conserved 27-amino acid GroEL fragment and docked it against the Aβ (1-42) tetramer. Interaction analysis revealed stabilizing salt bridges, hydrogen bonds, and extensive non-bonded contacts within the GroEL–Aβ complex. Molecular dynamics simulations (50 ns) demonstrated that GroEL binding enhanced Aβ oligomer stability, evidenced by reduced structural deviations and a more extensive hydrogen bonding network compared to Aβ oligomer alone. These computational findings support a novel mechanism whereby *H. pylori* GroEL directly stabilizes soluble Aβ oligomers. We hypothesize that this stabilization inhibits their aggregation into plaques while paradoxically prolonging the lifetime of neurotoxic species, potentially increasing neurodegeneration through pathways distinct from canonical amyloid deposition. This highlights the complex role of bacterial proteins in AD and underscores the need for experimental validation of GroEL–Aβ interactions as a potential therapeutic target.

## 1. INTRODUCTION

Alzheimer’s disease (AD) is the most common cause of dementia in the world, and mostly common in the elderly population (Mayeux and Stern 2012). It is neuropathologically characterized by extracellular accumulation of amyloid-beta (Aβ) in plaques and intracellular neurofibrillary tangles of hyperphosphorylated tau protein (Serrano-Pozo et al. 2011). According to the amyloid cascade hypothesis, the accumulation of Aβ is the main cause of AD pathogenesis. Notably, recent research findings suggest that soluble Aβ oligomers, rather than insoluble plaques, are the primary neurotoxic species that cause memory loss and synaptic impairment in AD (Goure et al. 2014).

Although ageing and genetics are significant risk factors, chronic microbial infections and inflammation are emerging as potential causes of AD (Hersi et al. 2017; Albaret et al. 2020). *Helicobacter pylori* (*H. pylori*) is a widespread gastric pathogen that is linked to extra-gastric diseases, such as neurodegenerative disorders. *H. pylori* infection has the potential to cause systemic inflammation, disrupt the blood-brain barrier (BBB), and stimulate the mechanisms involved in AD pathology (Álvarez-Arellano 2014; Franceschi et al. 2015).

Notably, *H. pylori* releases outer membrane vesicles (OMVs), which transport virulence factors such as the chaperonin GroEL that enter the BBB and cause neuroinflammation (Palacios et al. 2023). GroEL is a bacterial homolog of human Hsp60, which plays a central role in the protein folding process and could control amyloidogenic pathways (Wälti et al. 2017). However, the role of bacterial chaperonins in AD appears complex. Recent studies prove that they can have neuroprotective properties by inhibiting the formation of Aβ fibrils (Wälti et al. 2018) and other studies show that they can activate neurodegeneration (Palacios et al. 2023). This paradox highlights a critical gap in how bacterial chaperonins modulate the stability and toxicity of intermediate Aβ aggregates.

This study hypothesizes that *H. pylori* GroEL binds to and stabilizes soluble Aβ oligomers, an interaction that could potentiate the lifetime and toxicity of these species at neuronal membranes, thereby influencing AD progression through pathways distinct from plaque accumulation. To examine this, a computational methodology was used, which comprised pan-genome analysis of *H. pylori* strains, functional annotation, structural modeling, protein-protein docking, and molecular dynamics simulations. This will be used to explain GroEL conservation across strains as well as for mapping its molecular interactions with Aβ oligomers at the molecular level, which can be used to determine the microbial role in AD pathogenesis and identify potential therapeutic strategies.

## 2. METHODOLOGY

### 2.1. Genomic Data Acquisition and Curation

A comprehensive dataset of *Helicobacter pylori* genomes was compiled to investigate the conservation and functional role of the GroEL (Cpn60) protein. The National Center of Biotechnology (NCBI) genome database was used to download 353 complete *H. pylori* genome sequences on September 25, 2024 (Sayers et al. 2011). Complete assemblies were used to ensure high-quality and consistency of data to use in downstream analysis.

### 2.2. Genome Quality Assessment and Annotation

The genome assemblies were quality assessed by the QUAST tool (Quality Assessment Tool for Genome Assemblies, version 5.3.0) (Gurevich et al. 2013) on the Galaxy platform (Afgan et al. 2018). Subsequent genome annotation was performed using Prokka (version 1.14.6) (Seemann 2014) within the Galaxy platform. The pan-genome analysis was done using the generated GFF3 files to provide a common annotation framework to all genomes.

### 2.3. Pan-Genome Analysis

The pan-genome was analyzed, and variation and conservation of gene content were examined among the 353 *H. pylori* strains. The annotated GFF3 files were processed using Roary (version 3.13.0) (Page et al. 2015), a tool designed for rapid and scalable pan-genome construction. Roary clustered genes into core (present in ≥99% strains), soft core (95-99%), shell (15-95%), and cloud (<15%) categories, enabling characterization of the genomic diversity. Key outputs included a gene presence-absence matrix and summary statistics defining the distribution of gene categories. The gene encoding GroEL was specifically identified within the matrix for conservation analysis across strains. The pan-genome dynamics were further visualized with pan-core genome plots, illustrating core gene stability and pan-genome growth, while Heap’s law fitting quantified the openness of the genome.

### 2.4. Functional Annotation

Functional characterization of pan-genome proteins was performed using InterProScan (version 5.59-91.0) (Zdobnov and Apweiler 2001; Quevillon et al. 2005; Hunter et al. 2009; Jones et al. 2014) on the Galaxy platform. The large protein cluster dataset generated by Roary was split into manageable subsets and submitted for domain and motif analysis against integrated databases InterProScan such as Pfams, SMART, and PROSITE. Results included protein families, conserved domains, and Gene Ontology (GO) annotations. GroEL identification was refined by filtering for InterPro accession IPR001844 (Chaperonin Cpn60/GroEL) and related signatures. To target the functionally significant region, sequences annotated with GO terms related to protein folding and ATP-dependent protein refolding (GO:0042026, GO:0140662) were extracted for downstream structural analysis.

### 2.5. Structural Modelling of GroEL Fragment

A 27-amino acid conserved fragment of GroEL (VKVTMGPRGRNVLIQKSYGAPSITKDG) corresponding to a region of high sequence conservation and functional relevance was selected for structural modelling. Due to the absence of experimentally determined structures for this fragment, computational prediction was carried out using AlphaFold3 (Jumper et al. 2021; Abramson et al. 2024). Five candidate models were generated and assessed using average predicted Local Distance Difference Test score (pLDDT), predicted aligned error (PAE), ProSA Z-score (Wiederstein and Sippl 2007), Ramachandran plot analysis, and clashscore (Williams et al. 2018). The model with the highest structural reliability was chosen for subsequent molecular docking.

### 2.6. Molecular Docking Studies

The GroEL and amyloid beta (Aβ) oligomers interaction was investigated through protein-protein docking. The Aβ (1-42) tetramer (PDB ID: 6RHY) (Ciudad et al. 2020) was obtained from the Protein Data Bank (Berman et al. 2000). The fragment of GroEL was docked against the Aβ tetramer through the HDOCK server (Yan et al. 2017). Blind docking was used to enable full, impartial sampling of the Aβ. To determine chain-specific binding, the interface between the GroEL fragment and each of the individual Aβ chains (A, B, C, D) in the tetramer was measured individually. In each of the simulations, a set of poses was created and sorted by docking score and confidence score. The most ranked pose was chosen to be analyzed further. Analysis after docking comprised visualizing docked complexes with the help of Chimera (Pettersen et al. 2004), and residues interface mapping and interaction characterization analysis were performed using PDBsum (Almo et al. 1997).

### 2.7. Protein-Protein Interaction Analysis

The top docked GroEL–amyloid beta complex was subjected to rigorous interaction analysis using PDBsum (Almo et al. 1997), focusing on interface residues, area, electrostatic interactions, hydrogen bonds, and non-bonded contacts. Criteria for salt bridges included charged residues within 3.2 Å, hydrogen bonds included donor-acceptor distances <3.5 Å with angles ≥120°, and van der Waals contacts considered within 4.0 Å. This detailed interaction mapping identified stabilizing contacts, critical residues, and the physicochemical basis of complex stability.

### 2.8. Molecular Dynamics Simulations

Molecular dynamics simulations were conducted using the UAMS SimLab platform (https://simlab.uams.edu/) (Abraham et al. 2015) to evaluate the stability of the GroEL–amyloid beta complexes. The top docked complex was prepared with the CHARMM36 force field and solvated in a cubic box of explicit TIP4P water molecules, maintaining a minimum 10 Å buffer around the protein. The system was neutralized with 0.15 M NaCl. The steepest descent algorithm was applied to minimize the energy, and the subsequent 50,000 steps were run until the equilibrations were achieved in NVT and NPT ensembles at 310 K and 1 bar, respectively. The production runs were performed by 50 ns, with 10 ps coordinates being recorded.

The docked complexes of the GroEL with each of the individual amyloid beta chains were considered as isolated protein molecules and subjected to protein-in-water molecular dynamics simulations. Upon analysis, the GroEL–amyloid beta chain B complex exhibited the most favorable stability profile. Consequently, protein-in-water simulations of amyloid beta chain B alone were also performed for comparison. Trajectory analyses, including Root Mean Square Deviation (RMSD), Root Mean Square Fluctuation (RMSF), Solvent Accessible Surface Area (SASA), and hydrogen bonding, were then conducted to compare the unbound peptide with the GroEL-bound complex.

## 3. RESULTS

### 3.1. Genome Dataset, Quality Assessment, and Annotation

Quality assessment of the curated dataset of 353 complete *H. pylori* genomes confirmed the assemblies were nearly complete with an average of 1.33 contigs per genome, genome sizes approximating the expected ∼1.6 Mbp, and GC content averaging 38.9%. These metrics confirmed low fragmentation and high reliability of the dataset. Genome annotation by Prokka identified coding sequences, tRNAs, rRNAs, and other features, providing a uniform basis for subsequent pan-genome and comparative analyses.

### 3.2. Pan Genome Analysis

Pan-genome analysis of the 353 *H. pylori* genomes identified a total of 17,217 gene clusters categorized by occurrence frequency across strains. Core genes, present in 99–100% of strains, numbered 643 and likely represent essential functions. Soft core (95–99%), shell (15–95%), and cloud (<15%) genes numbered 129, 1,238, and 15,207, respectively, indicating the large and dynamic accessory genome of *H. pylori* **(Figure 1A).**

**Figure 1.**
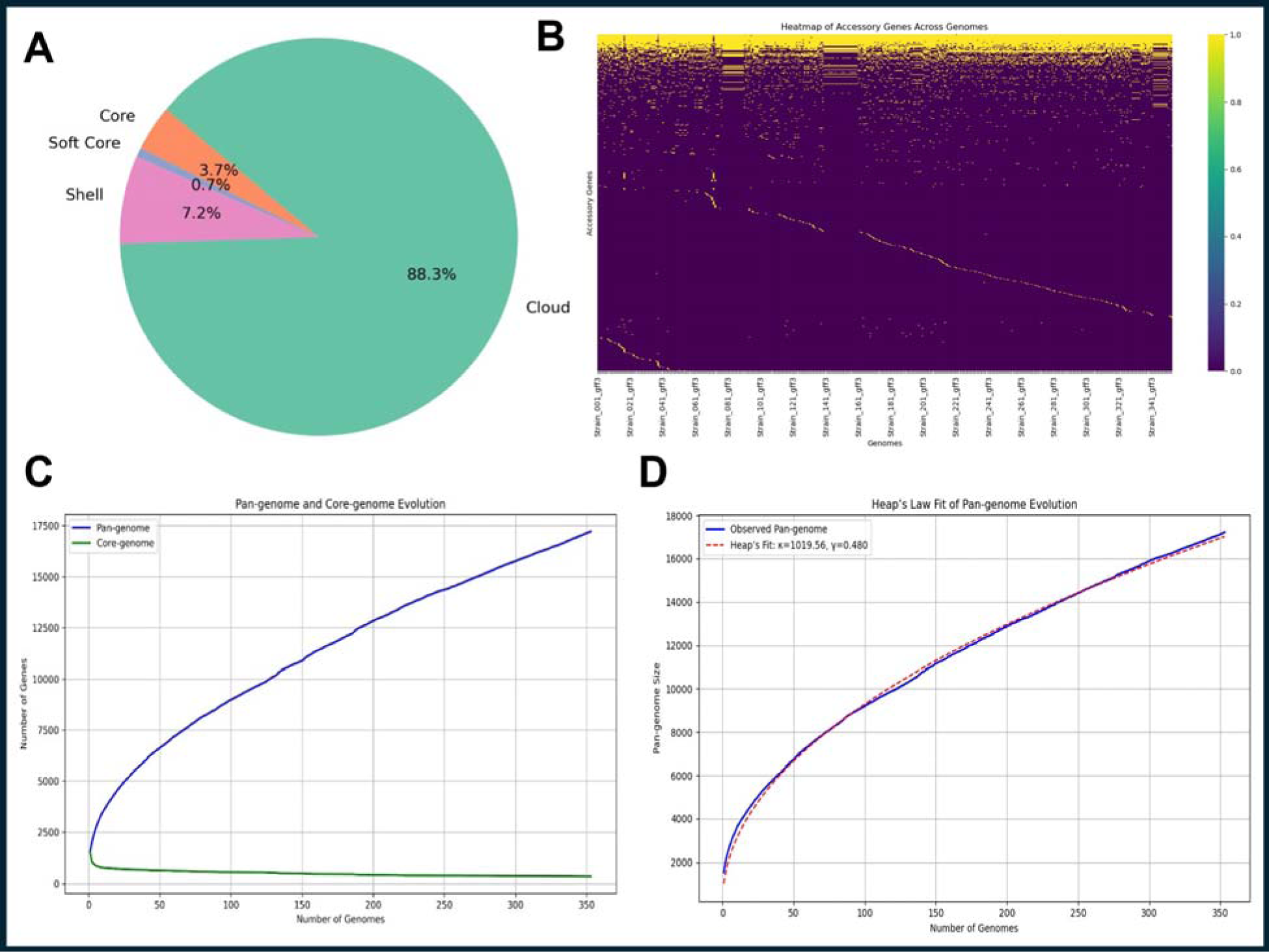
Comprehensive pan-genome analysis of 353 Helicobacter pylori genomes. **A.** Pie chart showing the distribution of core (643), soft core (129), shell (1,238), and cloud (15,207) genes in the H. pylori pan-genome. **B.** Heatmap displaying the presence (colored) and absence (contrasting) of accessory genes across 353 genomes, illustrating gene sharing patterns and diversity. **C.** Pan-core genome plot showing a declining core gene curve (green line) and expanding pan-genome curve (blue line), indicating an open pan-genome. **D.** Heap’s law fit showing the expansion of the H. pylori pan-genome; the sublinear trend (γ < 1) confirms its openness and ongoing gene acquisition.

The small core genome relative to the expansive accessory genome reflects the extensive genetic diversity and adaptability of *H. pylori*. Analysis of the pan-core genome plot revealed a declining core gene count and a steady expansion of the pan-genome with each additional genome **(Figure 1C)**, indicative of an open pan-genome. This was quantitatively supported by Heap’s law fitting **(Figure 1D)**, confirming ongoing gene acquisition among strains. Accessory genome heatmaps revealed clustering of strains with similar accessory gene profiles, reflecting evolutionary and ecological diversity **(Figure 1B).** These findings highlight the genetic plasticity and niche adaptation of *H. pylori*, as well as the ongoing discovery of novel genes through expanded sequencing efforts.

### 3.3. Functional Annotation of the *H. pylori* Pan-Genome

Functional annotation of clustered proteins derived from the pan-genome was performed using InterProScan, integrating multiple protein signature databases to identify conserved domains and assign Gene Ontology (GO) terms. Given the large dataset, proteins were processed in manageable subsets to ensure comprehensive annotation coverage. Annotation results revealed diverse protein families and functional domains across the pan-genome.

Focusing on the chaperonin GroEL (Cpn60), it was found to be highly conserved, present in 293 out of 353 strains (83%), highlighting its essential role in *H. pylori* biology. Multiple domain signatures and GO terms related to protein folding and ATP-dependent refolding (GO:0042026, GO:0140662) were consistently observed across strains **(Table S1).** Based on this, a 27-amino acid conserved region of GroEL (VKVTMGPRGRNVLIQKSYGAPSITKDG) implicated in chaperone function was selected for detailed structural and interaction analyses **(Table 1).**

**Table 1.** Conserved GroEL fragment selected for docking.

| Protein Accession | Sequence Fragment | GO Terms | InterPro Annotation |
| --- | --- | --- | --- |
| groL_Strain_066_00981 | **VKVTMGPRGRNVLIQ**<br>**KSYGAPSITKDG** | GO:0042026,<br>GO:0140662 | IPR001844<br>(Chaperonin<br>Cpn60) |

### 3.4. Structural Modeling and Quality Assessment of the GroEL Fragment

The conserved 27-amino acid GroEL fragment (VKVTMGPRGRNVLIQKSYGAPSITKDG) was modeled using AlphaFold3, producing five candidate 3D structures. Model quality was assessed quantitatively using average pLDDT, predicted aligned error (PAE), ProSA Z-score, Ramachandran plot statistics, and clashscore **(Table S2).** Despite uniform low confidence coloring in the pLDDT, likely due to the fragment’s short length, Model 1 was selected for further analysis based on its superior metrics, including the highest average pLDDT and lowest predicted error. The selected model provides a reliable structural basis to investigate molecular interactions of GroEL with amyloid beta oligomers through docking and simulation studies.

### 3.5. Docking Analysis of GroEL Fragment with Amyloid-Beta Oligomer

The conserved GroEL fragment was docked against each chain (A, B, C, D) of the amyloid beta (1-42) tetramer (PDB ID: 6RHY) using the HDOCK server. For each chain, 100 docking poses were generated and ranked by docking and confidence scores **(Table 2).** The top-ranked model for chain B exhibited the most favorable docking score (−176.56) and the highest confidence score (0.6062), indicating a stable binding conformation. The best 10 docking and confidence scores of A, B, C and D chain complexes are mentioned in **Tables S3-S6.** Visualization of the best docking poses of each chain, especially chain B, revealed consistent interaction patterns between the GroEL fragment **(Figures 2A-2E).** The best docking pose of this complex was selected for detailed protein-protein interaction and molecular dynamics analyses.

**Figure 2.**
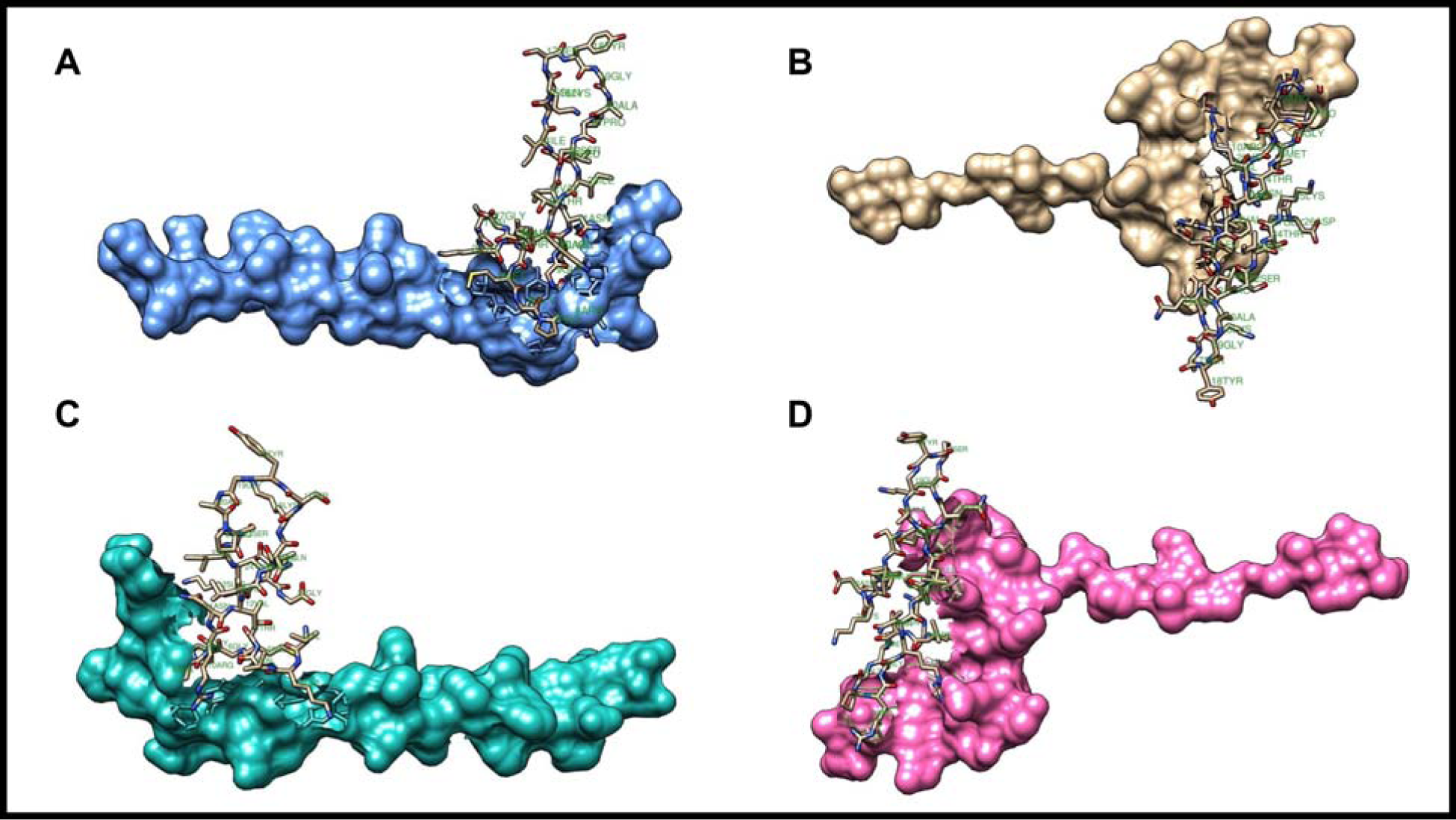
Best docked models of GroEL fragment with amyloid beta chains A–D. Representative docked complexes of the GroEL fragment with each Aβ chain (A–D) are visualized in Chimera. Amyloid-β chains are displayed as surface models with distinct colors: **A.** chain A in cornflower blue, **B.** chain B in gold, **C.** chain C in light sea green, and **D.** chain D in hot pink. The GroEL fragment is shown as a stick model in all panels, highlighting the protein– protein interaction interface with each Aβ chain.

**Table 2.**
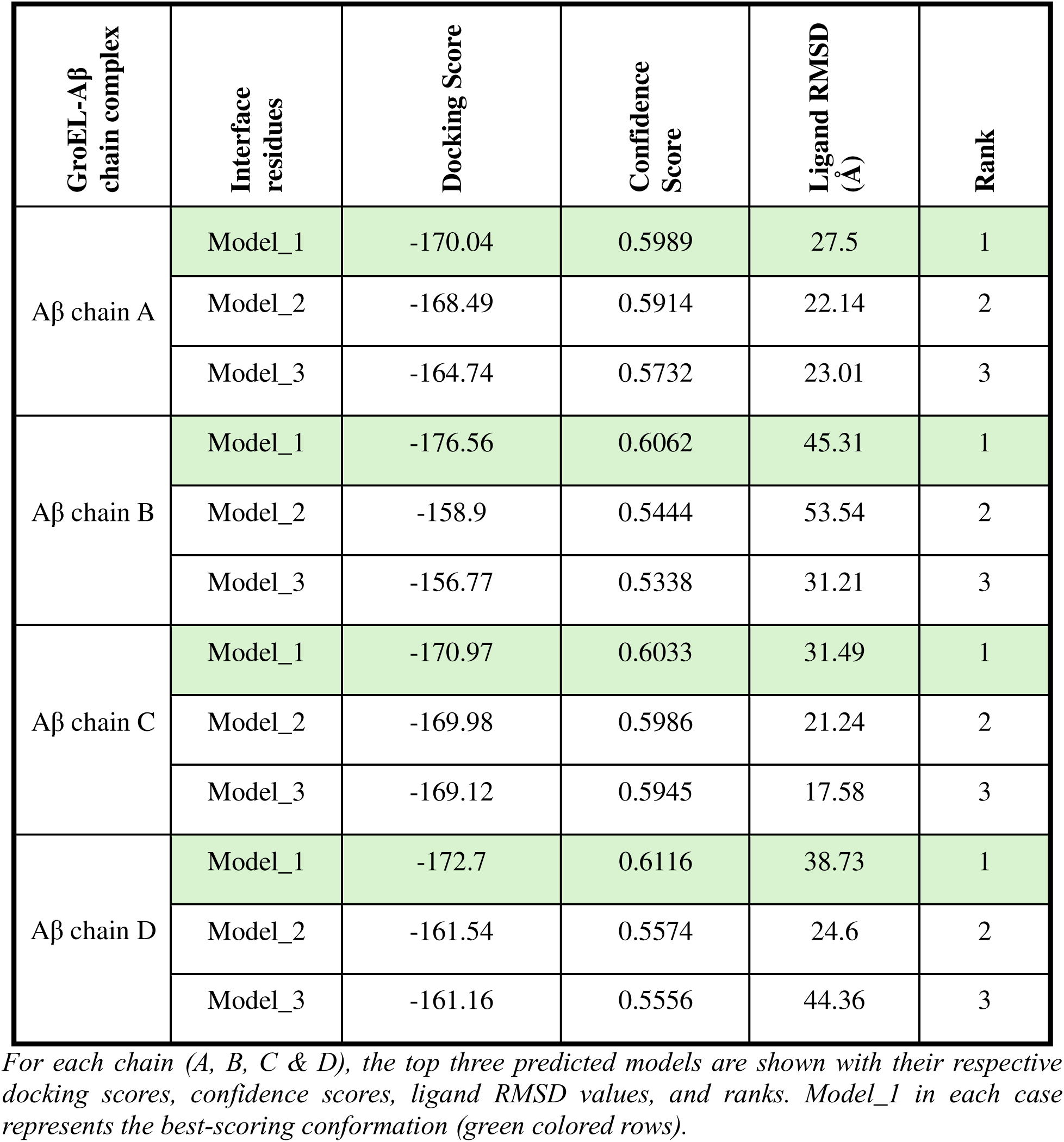
HDOCK results of each chain of the amyloid beta complexes with GroEL.

### 3.6. Protein–Protein Interaction Analysis of the Selected Docked Complex

The highest-ranked docked complex between the conserved GroEL fragment and amyloid bet chain B was subjected to detailed interface analysis using PDBsum. The interaction interface involved 14 residues of GroEL and 11 residues of amyloid beta, covering an interface area of approximately 564–669 Å² **(Table 3).** The complex was stabilized by three salt bridges (Arg8– Glu3, Arg10–Asp7, Arg10–Glu11), one hydrogen bond (Arg10–Glu11), and around 100 non-bonded contacts indicative of van der Waals interactions **(Figure 3A-3B).** Ramachandran plot demonstrated good stereochemical quality, validating the structural reliability for further molecular dynamics simulations.

**Figure 3.**
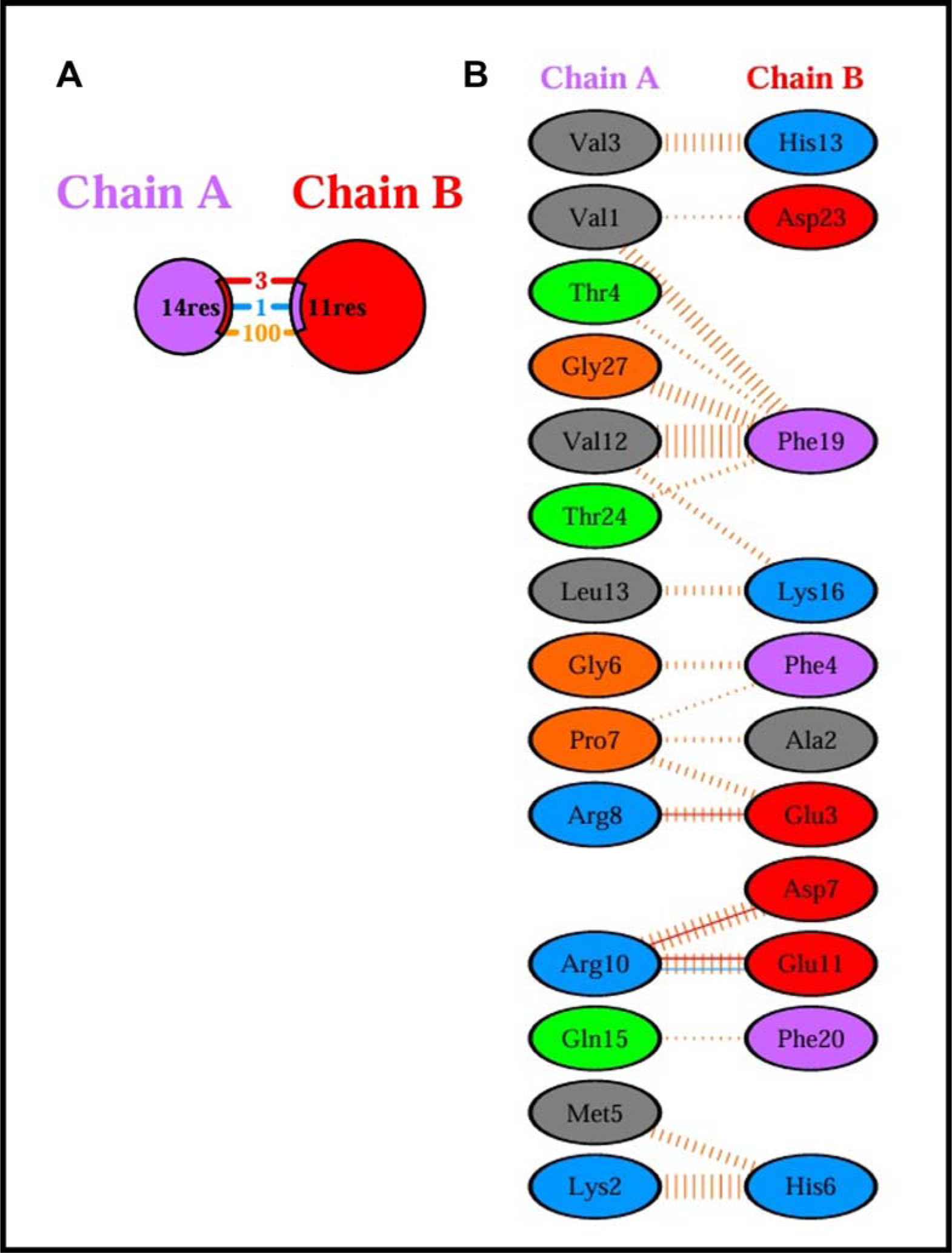
PDBsum analysis of the GroEL–amyloid beta chain B complex. **A.** Schematic representation of protein-protein interface obtained by PDBsum. **B.** Residue-level interaction map across the interface obtained from PDBsum. Residue-level interaction map highlighting 3 salt bridges (Arg8–Glu3, Arg10–Asp7, Arg10–Glu11), 1 hydrogen bond (Arg10–Glu11), and 100 non-bonded contacts between cpn60 and Aβ chain B.

**Table 3.** PDBsum interface statistics of the GroEL–amyloid beta chain B complex.

| **Chain** | **No. of Interface Residues** | **Interface Area (Å<sup>2</sup>)** | **No. of Salt Bridges** | **No. of Disulfide Bonds** | **No. of Hydrogen Bonds** | **No. of Non-bonded Contacts** |
| --- | --- | --- | --- | --- | --- | --- |
| GroEL (chain A) | 14 | 564 | 3 | N/A | 1 | 100 |
| Amyloid Beta chain B | 11 | 669 |  |  |  |  |

Here 'N/A' indicates 'Not-applicable'

### 3.7. Molecular Dynamics Simulations of GroEL–Amyloid Beta Complexes

Molecular dynamics (MD) simulations of the GroEL fragment complexed with amyloid-β (Aβ) chains A, B, C, and D were performed for 50 ns to assess their structural stability and interaction dynamics **(Figure 4).** All complexes demonstrated stable trajectories; among the four, the GroEL–Aβ chain B complex exhibited the highest structural stability, maintaining the lowest RMSD (< 1.5 nm) and minimal fluctuations, while chains A and C displayed transient conformational shifts with RMSD peaks reaching approximately 4.0 nm and 3.4 nm, respectively **(Figure 4A).** RMSF profiles localized most flexibility to solvent-exposed or terminal regions, with chain B showing the lowest average flexibility (0.75 nm) and chain C the highest (1.1 nm) **(Figure 4B).**

**Figure 4.**
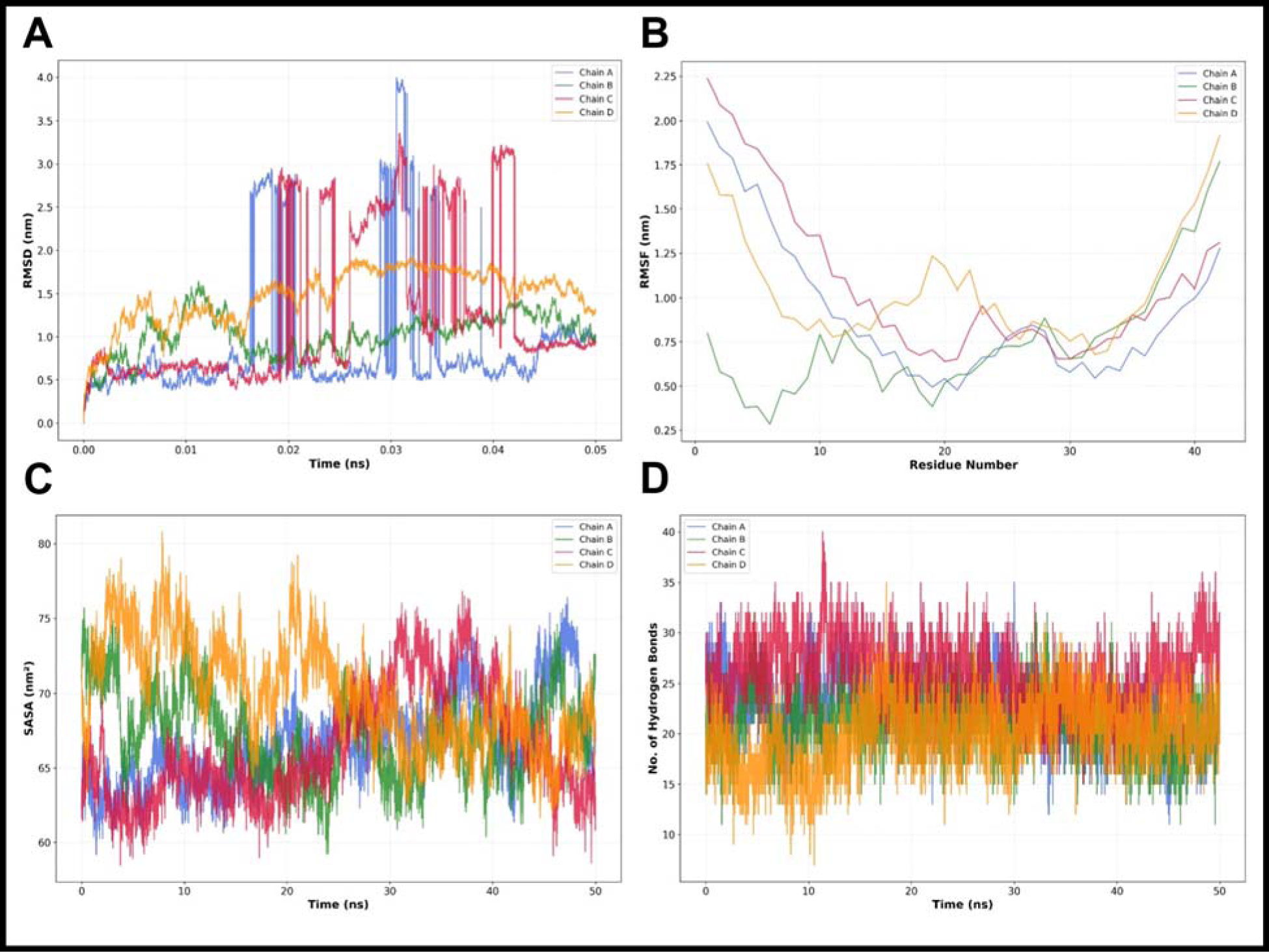
Comparative molecular dynamics (MD) analysis of GroEL complexes with amyloid-β (Aβ) chains A–D. **A.** RMSD plots showing that the GroEL–Aβ chain B (green) and chain D (orange) complexes exhibited the greatest structural stability, whereas chains A (blue) and C (pink) showed higher RMSD peaks (∼ 4.0 nm and 3.4 nm), indicating transient conformational shifts. **B.** RMSF profiles revealing the lowest flexibility for chain B (green, 0.75 nm) and highest for chain C (pink, 1.1 nm), with moderate fluctuations for chain A (blue, 0.9 nm) and chain D (orange, 1.05 nm). **C.** SASA plots showing chain D (orange) with the highest mean surface area (70.0 nm²), while chains A, B, and C (blue, green, pink) exhibited similar values (∼ 66–67 nm²). **D.** Hydrogen-bond analysis showing chain C (pink) formed the most extensive network (∼ 26 bonds), followed by chain A (∼ 23, blue), chain B (∼ 21, green), and chain D (∼ 20, orange), supporting overall complex stability.

SASA analysis revealed moderate solvent exposure across all complexes, with chain D showing the highest mean SASA (70.0 nm²) and chains A, B, and C exhibiting comparable values (approximately 66–67 nm²) **(Figure 4C).** Hydrogen-bond analysis further confirmed the dynamic stability of these complexes, with chain C forming the most extensive network (∼ 26 bonds), followed by chains A (∼ 23 bonds), B (∼ 21 bonds), and D (∼ 20 bonds) **(Figure 4D).** Collectively, these MD results indicate that while all GroEL–Aβ complexes maintained structural integrity, the chain B complex displayed the most favorable stability and compactness throughout the simulation, consistent with its superior docking score and interaction profile.

#### 3.7.1. Comparative Dynamics of Amyloid-Beta Chain B Alone versus Complex with GroEL

Comparative molecular dynamics of the amyloid beta chain B alone and in complex with GroEL over 50 ns revealed pronounced stability improvements upon GroEL binding. The complex exhibited significantly lower RMSD (mean 1.02 nm) than free chain B (mean 1.40 nm), reflecting reduced structural deviations **(Figure 5A, Table S7).** RMSF per residue analysis showed a slight global flexibility reduction in the complex (0.746 nm vs 0.762 nm), complemented by localized stabilization of 16 residues and increased flexibility in 26 others **(Figure 5B, Table S8).** Solvent accessible surface area markedly increased upon complex formation (67.2 nm² vs 45.7 nm²), suggesting GroEL reduces peptide compaction or aggregation **(Figure 5C, Table S9).** Hydrogen bonding analysis further highlighted an enhanced interaction network in the complex (mean 21.2 bonds) compared to the peptide alone (9.0 bonds), consistent with improved stability **(Figure 5D, Table S10).** The summary table of comparative MD simulation statistics for all the key parameters analysis is shown in **Table 4**.

**Figure 5.**
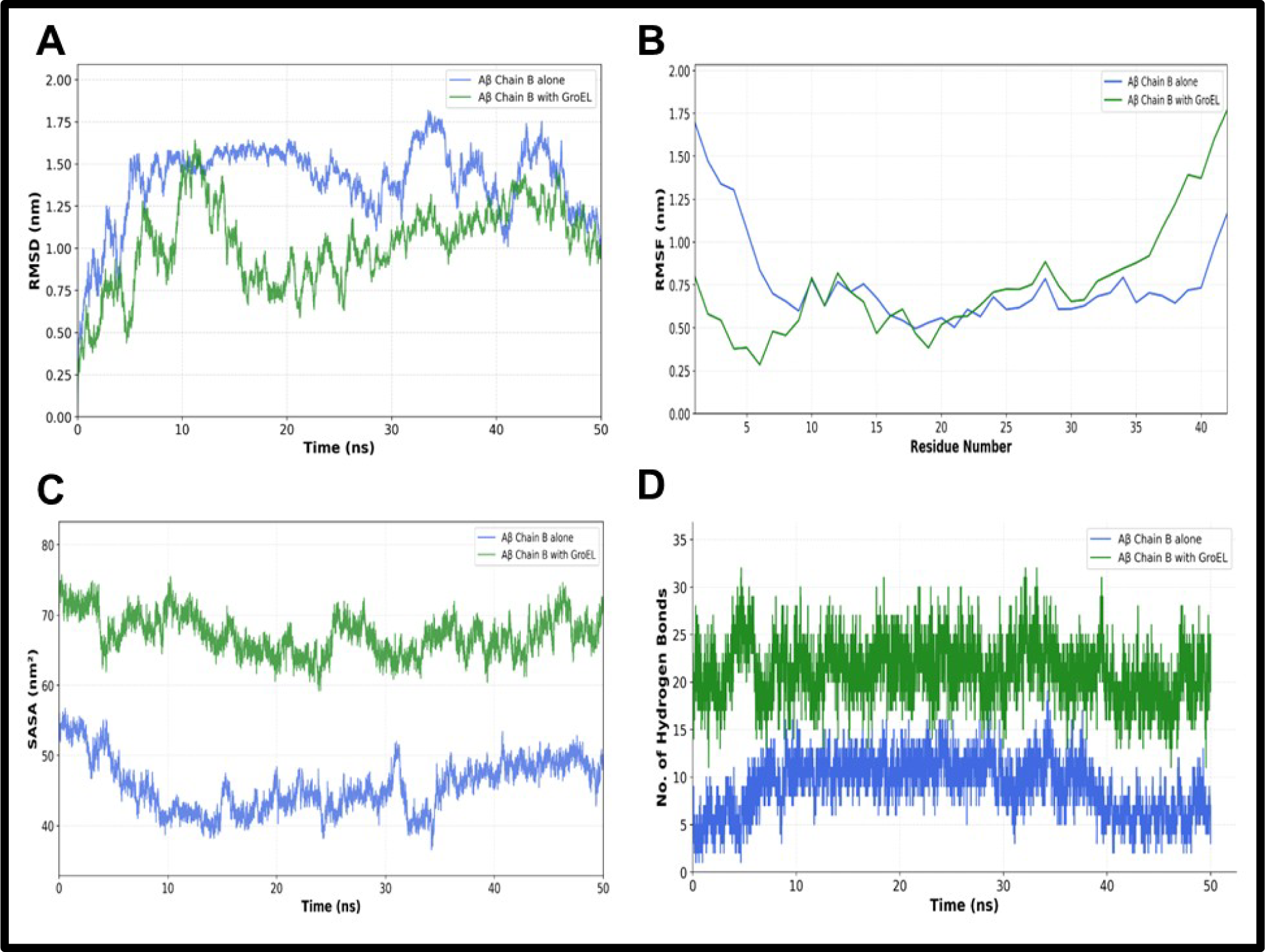
Comparative molecular dynamics (MD) analysis of amyloid-β (Aβ) chain B alone and in complex with H. pylori GroEL. **A.** RMSD plot showing reduced structural deviations in the GroEL–Aβ complex (green) compared to Aβ alone (blue), indicating higher stability. **B.** RMSF analysis showing decreased flexibility in the GroEL-bound complex (green) relative to Aβ alone (blue). **C.** Solvent Accessible Surface Area (SASA) profiles indicating increased surface exposure for the GroEL–Aβ complex (green), suggesting reduced compaction. **D.** Hydrogen bond analysis showing a higher number of stable H-bonds in the GroEL–Aβ complex (green) than in Aβ alone (blue), supporting enhanced structural stability.

**Table 4.** Summary table of comparative MD simulation statistics.

| **Parameter** | **Aβ Chain B Alone** | **Aβ Chain B with GroEL** |
| --- | --- | --- |
| Mean RMSD (nm) | 1.40 | 1.02 |
| Mean RMSF (nm) | 0.76 | 0.75 |
| Mean SASA (nm<sup>2</sup>) | 45.66 | 67.21 |
| Mean H-bonds | 9.01 | 21.23 |

## 4. DISCUSSION

This study provides computational evidence for a novel mechanism by which *H. pylori* may influence AD pathology, moving beyond its established role in neuroinflammation. Our combination of pan-genomes, structural modelling, and molecular dynamics shows that the bacterial chaperonin GroEL, a highly conserved aspect of *H. pylori* OMVs, is capable of binding and stabilizing the toxic soluble Aβ oligomers directly.

We verified the wide range of genetic diversity with our analysis of a high-quality dataset of 353 complete *H. pylori* genomes with a small core genome and an extensive and open pan-genome. This genomic plasticity promotes the adaptability of the bacterium and implies that effector proteins that could be shared among strains are of core significance (Ali et al. 2015; Uchiyama et al. 2016). In this context, GroEL (Cpn60) was found to be a highly conserved protein, which is found in 83% of the strains studied. It has a fundamental role in protein folding and cellular homeostasis (Hartl et al. 2011), thereby being an attractive prospect as a mediator of bacterium-host interaction, especially in the setting of protein misfolding diseases such as AD. Importantly, the recently conducted research showed that *H. pylori* OMVs supplemented with GroEL are capable of disrupting the BBB, which can result in neuroinflammation and, consequently, neurodegenerative mechanisms (Palacios et al. 2023).

Other studies have proposed a neuroprotective potential of bacterial chaperonins GroEL, which inhibits the development of Aβ fibrils, therefore slowing down the formation of plaques (Wälti et al. 2018). Our findings, on the contrary, develop a more subtle and possibly harmful hypothesis. We propose that *H. pylori* GroEL stabilizes toxic soluble Aβ oligomers, the most neurotoxic form in AD. Molecular docking revealed a strong and specific interaction between a conserved 27-amino acid GroEL fragment and the Aβ tetramer, with the most stable binding pose identified for chain B.

Molecular dynamics simulations provided direct evidence for this stabilization. The GroEL–Aβ complex exhibited a significantly lower RMSD than the Aβ oligomer alone, indicating enhanced global structural stability. This was accompanied by a substantial increase in SASA and hydrogen bonding network. While the overall residue flexibility (RMSF) changed only modestly, localized variations indicated a subtle remodelling of the oligomer’s dynamics, with specific residues rigidifying at the interaction interface.

Therefore, the seemingly paradoxical roles of GroEL, such as inhibiting fibril formation while stabilizing soluble oligomers, reflect the complexity of amyloid pathology and bacterial protein interactions in AD. While delaying fibril formation may offer neuroprotection by preventing plaque deposition, stabilization of soluble oligomers could increase neurotoxicity, underscoring the need for further experimental investigation. This work contributes molecular-level insight into how *H. pylori* influences AD pathology through GroEL beyond classical inflammatory pathways, highlighting a potential microbial modulator of protein misfolding diseases such as AD.

Despite these significant findings, several limitations must be acknowledged. These results are based primarily on in silico analyses, and although computational approaches provide valuable mechanistic insights, biological validation through in vitro and in vivo studies is important. Additionally, focusing on a conserved GroEL fragment may not capture the full spectrum of interactions mediated by the complete chaperonin or other bacterial proteins within OMVs. Future research should expand experimental validation, explore the functional consequences of GroEL-Aβ interactions on neuronal health, and consider microbial contributions to AD in clinical contexts.

In conclusion, this integrated pan-genomic, structural, and dynamic study presents strong computational evidence that *H. pylori* GroEL stabilizes soluble, neurotoxic Aβ oligomers, potentially influencing AD progression through mechanisms distinct from plaque inhibition. This novel insight enriches the growing understanding of microbial roles in neurodegeneration and could open avenues for targeted therapeutic interventions aimed at modulating bacterial protein interactions with host amyloidogenic pathways.

## 5. CONCLUSION

This paper provides a comprehensive computational analysis of the genomic diversity of *H. pylori* and molecular interactions of the conserved chaperonin GroEL with amyloid beta oligomers playing a role in the pathology of AD. Strict pan-genome investigation exposed a very adaptable *H. pylori* genome having a small core as well as a broad collection of accessory genes, emphasizing the adaptability of the species. Broad conservation of GroEL was validated by functional annotation, which has a focus on the critical role of this protein in biology. Structural modeling and protein-protein docking revealed specific binding of a conserved fragment of GroEL to amyloid beta oligomers, especially stabilizing the soluble and neurotoxic forms associated with the development of AD. Simulations using molecular dynamics showed that the increase in the stability of the complexes caused by lower RMSD and RMSF values, higher SASA, and an expanded hydrogen-bonding network compared to Aβ alone.

These results confirm a new way in which *H. pylori* GroEL could regulate AD pathology through stabilization of soluble Aβ oligomers, which provides new insights into the role of microbes in neurodegeneration. Although this is a computational study, it provides sufficient groundwork for the future use of experimental validation and discusses the possible therapeutic intervention of GroEL-Aβ interactions. Integrating microbial genomics, bioinformatics, and molecular modeling in this context increases the knowledge of the host-microbe interaction in neurodegenerative diseases and opens up interdisciplinary studies that involve microbiology, neurobiology, and computational biology.

## DECLARATIONS

## AUTHOR CONTRIBUTIONS

**W.M.A.** performed all computational experiments, including database curation, pangenome analysis, protein modelling, docking, and MD simulation analysis. Also, made the figures and prepared the manuscript. Mr. **I.A.** supervised the work and reviewed the manuscript.

## DATA AVAILABILITY

Not applicable

## CODE AVAILABILITY

Not applicable

## Abbreviations

(AD): Alzheimer’s disease
(Aβ): Amyloid beta
(H. pylori): Helicobacter pylori
(GO): Gene Ontology
(MD): Molecular dynamics
(RMSD): Root mean square deviation
(RMSF): Root mean square fluctuation
(SASA): Solvent Accessible Surface Area

