## Supplementary Information for "Integrative Computational Analysis Reveals *H. pylori* GroEL as a Stabilizer of Neurotoxic Amyloid-β Oligomers"

*Table S1. Functional annotation snapshot of GroEL in strain 066*

| Protein Accession | Sequence Length | Analysis | Start Location | Stop Location | InterPro Accession | InterPro Description | GO Annotations |
|---|---|---|---|---|---|---|---|
| groL_Strain_066_00981 | 546 | FunFam | 191 | 372 | - | - | - |
| groL_Strain_066_00981 | 546 | SUPERFAMILY | 1 | 229 | IPR027413 | GroEL-like equatorial domain superfamily | - |
| groL_Strain_066_00981 | 546 | Gene3D | 5 | 518 | IPR027413 | GroEL-like equatorial domain superfamily | - |
| groL_Strain_066_00981 | 546 | Gene3D | 136 | 409 | IPR027410 | TCP-1-like chaperonin intermediate domain superfamily | - |
| groL_Strain_066_00981 | 546 | Gene3D | 191 | 372 | IPR027409 | GroEL-like apical domain superfamily | - |
| groL_Strain_066_00981 | 546 | SUPERFAMILY | 316 | 521 | IPR027413 | GroEL-like equatorial domain superfamily | - |
| groL_Strain_066_00981 | 546 | PANTHER | 2 | 531 | - | - | - |
| groL_Strain_066_00981 | 546 | Hamap | 1 | 545 | IPR001844 | Chaperonin Cpn60/GroEL | GO:0042026, GO:0140662 |
| groL_Strain_066_00981 | 546 | ProSitePatterns | 404 | 415 | IPR018370 | Chaperonin Cpn60, conserved site | GO:0005524, GO:0006457 |
| groL_Strain_066_00981 | 546 | PRINTS | 267 | 290 | IPR001844 | Chaperonin Cpn60/GroEL | GO:0042026, GO:0140662 |
| groL_Strain_066_00981 | 546 | PRINTS | 397 | 418 | IPR001844 | Chaperonin Cpn60/GroEL | GO:0042026, GO:0140662 |
| groL_Strain_066_00981 | 546 | PRINTS | 82 | 109 | IPR001844 | Chaperonin Cpn60/GroEL | GO:0042026, GO:0140662 |
| groL_Strain_066_00981 | 546 | PRINTS | 26 | 52 | IPR001844 | Chaperonin Cpn60/GroEL | GO:0042026, GO:0140662 |
| groL_Strain_066_00981 | 546 | PRINTS | 349 | 374 | IPR001844 | Chaperonin Cpn60/GroEL | GO:0042026, GO:0140662 |
| groL_Strain_066_00981 | 546 | CDD | 3 | 520 | IPR001844 | Chaperonin Cpn60/GroEL | GO:0042026, GO:0140662 |
| groL_Strain_066_00981 | 546 | SUPERFAMILY | 183 | 375 | IPR027409 | GroEL-like apical domain superfamily | - |
| groL_Strain_066_00981 | 546 | TIGRFAM | 2 | 524 | IPR001844 | Chaperonin Cpn60/GroEL | GO:0042026, GO:0140662 |
| groL_Strain_066_00981 | 546 | Pfam | 22 | 520 | IPR002423 | Chaperonin Cpn60/GroEL/TCP-1 family | GO:0005524 |

*Here '-' indicates 'Not-determined'. The coloured row (green) represents the conserved GroEL fragment selected for docking.*

***Table S2. Quality assessment of AlphaFold models for the GroEL Fragment***

| Model | Avg pLDDT | Avg PAE | ProSA Z-score | Favoured Ramachandran (%) | Clashscore |
|---|---|---|---|---|---|
| 1 | 91.25 | 3.72 | 0.05 | 100% | 9.55 |
| 2 | 91.72 | 3.63 | 0.08 | 100% | 9.55 |
| 3 | 90.37 | 4.03 | 0.05 | 100% | 11.93 |
| 4 | 84.13 | 5.67 | 0.10 | 92% | 9.55 |
| 5 | 84.57 | 5.23 | -0.31 | 100% | 19.09 |

*The coloured row consists of the best model metrics.*

***Table S3. HDOCK results of amyloid beta chain A complexes with GroEL***

| Interface residues | Docking Score | Confidence Score | Ligand RMSD (Å) | Rank |
|---|---|---|---|---|
| Model_1 | -170.04 | 0.5989 | 27.50 | 1 |
| Model_2 | -168.49 | 0.5914 | 22.14 | 2 |
| Model_3 | -164.74 | 0.5732 | 23.01 | 3 |
| Model_4 | -164.13 | 0.5702 | 17.11 | 4 |
| Model_5 | -160.86 | 0.5541 | 26.07 | 5 |
| Model_6 | -159.44 | 0.5471 | 24.17 | 6 |
| Model_7 | -159.08 | 0.5453 | 20.34 | 7 |
| Model_8 | -158.53 | 0.5425 | 16.46 | 8 |
| Model_9 | -158.51 | 0.5424 | 14.31 | 9 |
| Model_10 | -156.48 | 0.5324 | 23.09 | 10 |

*The coloured row consists of the best model metrics.*

***Table S4. HDOCK results of amyloid beta chain B complexes with GroEL***

| **Interface residues** | **Docking Score** | **Confidence Score** | **Ligand RMSD (Å)** | **Rank** |
|---|---|---|---|---|
| Model_1 | -176.56 | 0.6062 | 45.31 | 1 |
| Model_2 | -158.90 | 0.5444 | 53.54 | 2 |
| Model_3 | -156.77 | 0.5338 | 31.21 | 3 |
| Model_4 | -156.26 | 0.5313 | 26.17 | 4 |
| Model_5 | -155.95 | 0.5297 | 20.31 | 5 |
| Model_6 | -155.35 | 0.5267 | 33.07 | 6 |
| Model_7 | -155.16 | 0.5258 | 48.70 | 7 |
| Model_8 | -153.97 | 0.5198 | 26.13 | 8 |
| Model_9 | -153.74 | 0.5187 | 30.68 | 9 |
| Model_10 | -151.77 | 0.5088 | 27.53 | 10 |

*The coloured row consists of the best docking model metrics.*

*Table S5. HDOCK results of amyloid beta chain C complexes with GroEL*

| Interface residues | Docking Score | Confidence Score | Ligand RMSD (Å) | Rank |
|---|---|---|---|---|
| Model_1 | -170.97 | 0.6033 | 31.49 | 1 |
| Model_2 | -169.98 | 0.5986 | 21.24 | 2 |
| Model_3 | -169.12 | 0.5945 | 17.58 | 3 |
| Model_4 | -165.82 | 0.5784 | 29.17 | 4 |
| Model_5 | -165.16 | 0.5752 | 20.22 | 5 |
| Model_6 | -163.23 | 0.5658 | 19.47 | 6 |
| Model_7 | -159.80 | 0.5488 | 18.74 | 7 |
| Model_8 | -158.57 | 0.5427 | 26.76 | 8 |
| Model_9 | -157.17 | 0.5358 | 27.81 | 9 |
| Model_10 | -155.26 | 0.5263 | 30.97 | 10 |

*The coloured row consists of the best model metrics.*

*Table S6. HDOCK results of amyloid beta chain D complexes with GroEL*

| Interface residues | Docking Score | Confidence Score | Ligand RMSD (Å) | Rank |
|---|---|---|---|---|
| Model_1 | -172.70 | 0.6116 | 38.73 | 1 |
| Model_2 | -161.54 | 0.5574 | 24.60 | 2 |
| Model_3 | -161.16 | 0.5556 | 44.36 | 3 |
| Model_4 | -155.99 | 0.5299 | 21.64 | 4 |
| Model_5 | -154.96 | 0.5248 | 22.95 | 5 |
| Model_6 | -153.94 | 0.5197 | 44.17 | 6 |
| Model_7 | -153.75 | 0.5187 | 24.23 | 7 |
| Model_8 | -153.36 | 0.5168 | 23.65 | 8 |
| Model_9 | -153.33 | 0.5166 | 41.21 | 9 |
| Model_10 | -152.98 | 0.5149 | 24.05 | 10 |

*The coloured row consists of the best model metrics.*

**Table S7.** Comparative RMSD trajectory statistics of GroEL-Aβ chain B complex

| System | Mean RMSD (nm) | Std. Dev (nm) | Max RMSD (nm) |
|---|---|---|---|
| Aβ Chain B alone | 1.40 | 0.23 | 1.82 |
| Aβ-GroEL Complex | 1.02 | 0.25 | 1.64 |

**Table S*8*.** Comparative RMSF trajectory statistics of GroEL-Aβ chain B complex

| System | Mean RMSF (nm) | Std Dev (nm) | Max RMSF (nm) | Min RMSF (nm) | Residue Range |
|---|---|---|---|---|---|
| Aβ Chain B alone | 0.76 | 0.26 | 1.70 | 0.50 | 1–42 |
| Aβ-GroEL Complex | 0.75 | 0.32 | 1.77 | 0.28 | 1–42 (Aβ Chain B only) |

**Table S9.** Comparative SASA trajectory statistics of GroEL-Aβ chain B complex

| System | Mean SASA ($nm^2$) | Std Dev ($nm^2$) | Max SASA ($nm^2$) | Min SASA ($nm^2$) |
|---|---|---|---|---|
| Aβ Chain B alone | 45.66 | 3.74 | 56.71 | 36.57 |
| Aβ-GroEL Complex | 67.21 | 2.83 | 75.70 | 59.19 |

**Table S10.** Comparative hydrogen bond trajectory statistics of GroEL-Aβ chain B complex

| System | Mean H-bonds | Std Dev | Max | Min |
|---|---|---|---|---|
| Aβ Chain B alone | 9.01 | 2.93 | 19 | 1 |
| Aβ-GroEL Complex | 21.23 | 3.10 | 32 | 11 |
